# New proteomic biomarkers identified in plasma extracellular vesicles in sarcoidosis: a case-control matched study

**DOI:** 10.1101/2025.10.14.682425

**Authors:** Nan Miles Xi, Gabby Lea, Runzhen Zhao, Kamala Vanarsa, Mark Qiao, Dee Zhang, Jiwang Zhang, Chandra Mohan, Marc A. Judson, Laura L. Koth, Hong-Long Ji

**Author notes:** **To whom correspondence should be addressed:** Hong-Long (James) Ji, Nan Miles Xi (PhD),. Gabby Lea (PhD),. Runzhen Zhao (MD),. Kamala Vanarsa, (PhD),. Chandra Mohan (MD, PhD),. Marc A. Judson (MD),. Laura L. Koth (MD),. Hong-Long (James) Ji (MD, PhD),. Funding information: NIH HL134828, HL157533.

## Abstract

**BACKGROUND:** Sarcoidosis is a heterogeneous disease with unknown mechanisms, nonspecific therapies, and multiple etiologies. The role of blood extracellular vesicles (EVs) in the diagnosis and pathogenesis of sarcoidosis remains obscure. AIMS/OBJECTIVES. This study aims to test the hypothesis that the EV proteins in the blood can serve as phenotypic biomarkers of sarcoidosis. METHODS. We combined EV proteomics with machine learning algorithms to identify and prioritize biomarkers, enrich their functions, and cluster networks in case-control matched ACCESS patients. RESULTS. In total, 278 plasma EV proteins were significantly upregulated or downregulated in 40 sarcoidosis patients compared with 40 matched healthy controls. We identified 97 proteins that could serve as biomarkers with an AUC > 0.75. Of these, the AUC was > 0.90 for 13 proteins. 62 differentially expressed EV proteins strongly correlated with 20 clinical variables of severity, chest X-ray findings, and/or laboratory results. Functional annotation and network analysis suggest that these differentially expressed proteins regulate endocytosis, host responses to external stimuli, and transcription processes. Moreover, the top three ranked pathways were clathrin-mediated endocytosis, Hsp90 chaperone cycle, and spliceosome. CONCLUSIONS. This study demonstrates that plasma EV proteins can serve as biomarkers of various clinical phenotypes of the disease.

**At a Glance Commentary:** **Current Scientific Knowledge on the Subject:** Sarcoidosis is a heterogeneous condition affecting multiple organs. The role of blood extracellular vesicles in the diagnosis and pathogenesis of this condition remains unknown.

**What This Study Adds to the Field:** We identified differentially expressed proteins in plasma EVs by combining proteomics and machine learning algorithms. Top-ranked proteins can serve as diagnostic biomarkers and potential mechanisms for the development of sarcoidosis.

## Introduction

Inflammatory sarcoidosis is characterized by the formation of granulomas in involved tissues, with the chest being the most commonly affected (∼90%) ^1,2^. The severity of lung disease can be classified according to the Scadding stage. Besides the lungs, organs that are commonly involved with sarcoidosis include the skin, eyes, liver, lymph nodes, salivary glands, bones, joints, muscles, spleen, nervous system, kidneys, sinuses, and heart. Approximately 20% of patients are progressive and tend to develop lung fibrosis ^2–5^. The leading cause of death from sarcoidosis is respiratory failure associated with fibrotic lungs. The etiologies of sarcoidosis remain unclear despite its identification over a century ago. The prevalence, symptoms, potential triggers, and prognosis of the condition can vary significantly. Using traditional non-omics approaches (laboratory tests) and comparing statistical analysis, the following diagnostic biomarkers for sarcoidosis have been identified and validated: serum soluble interleukin-2 receptor (sIL-2R) ^6–13^, urinary U-8-OHdG ^14–16^, serum angiotensin-converting enzyme (ACE) ^6,7,11,12^, serum chitotriosidase ^17–19^, serum KL-6 ^20,21^, serum CRP ^11,21^, and serum BNP ^22,23^. IL12, IL18, MMP14, CTSS, amyloid A, ZNF688, ARFGAP1, CD14, LBP, α-2chain of haptoglobin, and PHA in serum and BAL may be useful biomarkers to distinguish sarcoidosis from healthy controls. In addition to serving as diagnostic biomarkers, some have also been suggested to predict lung function, inflammation, multiple organ involvement, organ failures, chronicity, and response to therapy. However, no study has evaluated the feasibility of identifying biomarkers in blood extracellular vesicles (EVs) for sarcoidosis. EVs can be isolated from as little as 5 microliters of human liquid samples^24^. Utilizing the purified EVs for omics studies offers the following advantages. First, the excessive plasma or serum proteins (e.g., albumin) can be removed without depleting the top 20 high-abundant proteins. Second, EV proteins exhibit long-term stability compared to bulk proteins, resulting in high detectability in EV proteins compared to cell-free plasma or serum ^25–29^. Third, EVs are a novel approach to identifying biomarkers, studying disease mechanisms, and clustering phenotypes of heterogeneous diseases ^26,27,30–38^. These studies combine advanced omics and machine learning algorithms. As recently reviewed, the feasibility of using EV proteomics as a strategy to study biomarker identification and molecular pathology in sarcoidosis remains an unanswered question ^39^.

This study aimed to identify differentiated biomarkers from healthy controls and their associations with clinical sarcoidosis phenotypes through the integration of high-throughput unbiased proteomics of plasma EVs and advanced machine learning algorithms. Our findings illustrate that, for the first time, novel biomarkers associated various phenotypes of the disease have been discovered through the analysis of ACCESS patients.

## Materials and Methods

### Patient cohorts

The participants were selected from the ACCESS (A Case Controlled Etiologic Study of Sarcoidosis) clinical trial ^40–44^. ACCESS was a case -control study of sarcoidosis where the clinical phenotype of sarcoidosis cases was described in detail and the control subjects were extremely well-matched to the cases. The cohort comprised of 40 sarcoidosis patients and 40 healthy controls matched for age (within 5 years) sex, race, and socioeconomic status/place of residence. The inclusion and exclusion criteria mirrored those of the ACCESS trial. The plasma samples and clinical dataset collected by the ACCESS trial were obtained through BioLINCC (Biologic Specimen and Data Repository Information Coordinating Center). The use of the plasma samples and de-identified clinical dataset was approved by the Institutional Review Board (IRB) of Loyola University Chicago (LU#216964). Patients were categorized based on their Scadding stage, organ involvement, chest radiographic findings, dyspnea score, pulmonary function, and blood tests. The collection, storage, and shipping of plasma samples were conducted according to the standard procedures recommended by the National Institutes of Health (NIH).

### Extraction of plasma extracellular vesicles (EVs)

EVs from plasma samples were captured and processed by Tymora Analytical Operations (West Lafayette, IN) using magnetic EVtrap beads as previously described ^45^. The isolated and dried EV samples were lysed to extract proteins using the phase-transfer surfactant (PTS)-aided procedure ^46^. The proteins were reduced and alkylated by incubating them in 10 mM TCEP and 40 mM CAA for 10 min at 95°C. The samples were then diluted fivefold with 50 mM triethylammonium bicarbonate and digested with Lys-C (Wako) at a 1:100 (wt/wt) enzyme-to-protein ratio for 3 h at 37°C. Trypsin was added at a final 1:50 (wt/wt) enzyme-to-protein ratio for overnight digestion at 37°C. To remove the PTS surfactants from the samples by acidification, trifluoroacetic acid (TFA) and an ethyl acetate solution were added to a final concentration of 1% TFA and at a 1:1 ratio, respectively. The mixture was vortexed for 2 min and then centrifuged at 16,000 × g for 2 min to separate the aqueous and organic phases. The organic phase (top layer) was removed, and the aqueous phase was collected. This step was repeated once more. The samples were dried in a vacuum centrifuge and desalted using Top-Tip C18 tips (Glygen) according to the manufacturer’s instructions. A portion of each sample was used to determine the peptide concentration using the Pierce Quantitative Colorimetric Peptide Assay. Finally, the samples were dried completely in a vacuum centrifuge and stored at -80°C.

### LC-MS/MS analysis

Each dried peptide sample was dissolved at 0.1 μg/μL in 0.05% trifluoroacetic acid with 3% (vol/vol) acetonitrile. 10 μL of each sample was injected into an Ultimate 3000 nano UHPLC system (Thermo Fisher Scientific). Peptides were captured on a 2-cm Acclaim PepMap trap column and separated on a heated 50-cm column packed with ReproSil Saphir 1.8 μm C18 beads. The mobile phase buffer consisted of 0.1% formic acid in ultrapure water (buffer A) with an eluting buffer of 0.1% formic acid in 80% (vol/vol) acetonitrile (buffer B) run with a linear 60-min gradient of 6–30% buffer B at a flow rate of 300 nL/min. The UHPLC was coupled online with a Q-Exactive HF-X mass spectrometer (Thermo Fisher Scientific). The mass spectrometer was operated in the data-dependent mode, in which a full-scan MS (from m/z 375 to 1,500 with a resolution of 60,000) was followed by MS/MS of the 15 most intense ions (30,000 resolution; normalized collision energy -28%; automatic gain control target (AGC) - 2E4, a maximum injection time - 200 ms; 60-sec exclusion).

### Data processing

The raw files were searched directly against the human UniProt database without redundant entries, using Byonic (Protein Metrics) and Sequest search engines loaded into Proteome Discoverer 2.3 software (Thermo Fisher Scientific). MS1 precursor mass tolerance was set at 10 ppm, and MS2 tolerance was set at 20 ppm. Search criteria included a static carbamidomethylation of cysteines (+57.0214 Da), variable modifications of oxidation (+15.9949 Da) on methionine residues, and acetylation (+42.011 Da) at the N-terminus of proteins. The search was performed with full trypsin/P digestion, allowing a maximum of two missed cleavages on the peptides analyzed from the sequence database. The false discovery rates (FDR) of proteins and peptides were set at 0.01. All protein and peptide identifications were grouped, and any redundant entries were removed. Unique peptides and unique master proteins were reported.

### Data preprocessing and profiling

The dataset of protein abundance generated from LC-MS contained a data matrix where each row represented a protein and each column a participant. Before statistical and machine learning analyses, the following preprocessing procedures were conducted on the data matrix to create a clean dataset: removing proteins that were not expressed in either the patient or control group, normalizing protein abundance through variance stabilization normalization (VSN) using the justvsn package in R ^47^, and imputing missing values using the random forest method with the R package missForest ^48^. The Wilcoxon rank sum test was used to compare the differences in abundance between patients and controls using the R function wilcox.test. The Benjamini-Hochberg (BH) method was utilized to control the FDR with the R function p.adjust. The fold change (FC) in the mean protein abundance between patients and controls was calculated. The differentially expressed proteins (DEP) were visualized in a heatmap with hierarchical clustering, volcano plot, and PCA plot using R packages ComplexHeatmap ^49^, EnhancedVolcano, and ggplot2 ^50^.

### Machine learning approaches for identifying biomarkers

To identify EV biomarkers, we trained a univariate logistic regression model using individual DEP to predict sarcoidosis patients and healthy controls. The logistic regression model was implemented using the R function glm. To assess the predictive performance of each protein, a five-fold cross-validation approach was repeated 100 times to calculate the area under the receiver operating characteristic (AUC), accuracy, sensitivity, specificity, positive predictive value (PPV), and negative predictive value (NPV). The probability threshold was set at 0.5. The differences in the protein expression levels of biomarker candidates between patients and controls were assessed using the Wilcoxon rank sum test. The visualization of AUC and protein expression was created using R package ggplot2 ^50^.

### Correlation analysis for the clinical relevance of biomarker candidates

To determine the clinical significance of the identified diagnostic biomarkers, their correlations with critical clinical variables were analyzed. The clinical variables include spirometry, Scadding stage, dyspnea score, blood tests, and chest X-ray readouts. Pearson correlation coefficients, p-values, and the slope of the linear regression line for continuous variables were calculated using the R function cor.test. The p-values were adjusted using the BH method to control the FDR. A significant correlation was considered if the adjusted p-value was < 0.05 or the Pearson correlation coefficient was > 0.4 and p-value < 0.05. The Wilcoxon rank sum test was conducted on the FC value of mean protein abundance to analyze the correlations between categorical variables and the identified biomarkers. The criteria for determining the significant correlations between biomarkers and categorical clinical variables were either an adjusted p-value < 0.05 or an absolute FC > 2.0 along with a p-value < 0.05. The linear regression was visualized using the geom_smooth function in the R package ggplot2 ^50^.

### Pathway enrichment and network analysis

As previously described ^51^, the R package pathfindR was used to enrich pathways for the identified biomarkers with an AUC > 0.750 ^52^. PathfindR considered the significance levels of individual biomarker proteins and utilized a protein-protein interaction network to enhance pathway enrichment outcomes. The run_pathfindR function in pathfindR package was executed to identify enriched pathways as defined in each database. To integrate the enriched pathways from different databases, pathway terms with an adjusted p-value of < 0.05 and at least two protein hits were included. The enriched pathways were hierarchically clustered by executing the cluster_enriched_terms function in the pathfindR package. The optimal number of clusters was determined by maximizing the average silhouette width. The pathway term with the lowest adjusted p-value in each cluster was selected as the representative pathway for that cluster and was included in the final enriched pathway result. The term_gene_graph function in pathfindR package was applied to create a network plot of the top-ranked pathways linked with associated biomarker proteins. The Heatmap function in the R package ComplexHeatmap was executed to perform hierarchical clustering of the biomarker proteins associated with the representative pathways ^49^.

### Determination of tissue and cellular origin

ToppGene was utilized to search the tissue and cellular origin of the DEPs ^53^. The hits of subcellular populations and tissues/organs were quantified and graphed as a function of their adjusted p-values.

### Sample size determination

The power analysis and calculation of the sample size were conducted using G power (version 3.1.9.7) ^54^. A two-tailed Wilcoxon-Mann-Whitney test was used to compare between-group differences in protein abundance and clinical variables. The expected effect size was set to 0.84 with a power of 95% and an FDR of 5%. Consequently, the sample size for sarcoidosis patients was set at 40 and for healthy controls at 40, to achieve the required power.

## Results

### 1. Baseline characteristics of participants

There were 80 participants (40 sarcoidosis cases and 40 controls) in the cohort (**Figure 1)**. There were no significant differences in demographic and baseline variables between the controls and the patient group (**Table 1**). The patients were represented at all Scadding stages (from stage 0 to stage 4). The predominant comorbidities included disorders of the heart, lungs, kidney, liver, rheumatologic system, neurologic system, endocrine system, and cancer. These comorbidities did not show significant differences, as eliminated by case-control matching and inclusion criteria.

**Figure 1.**
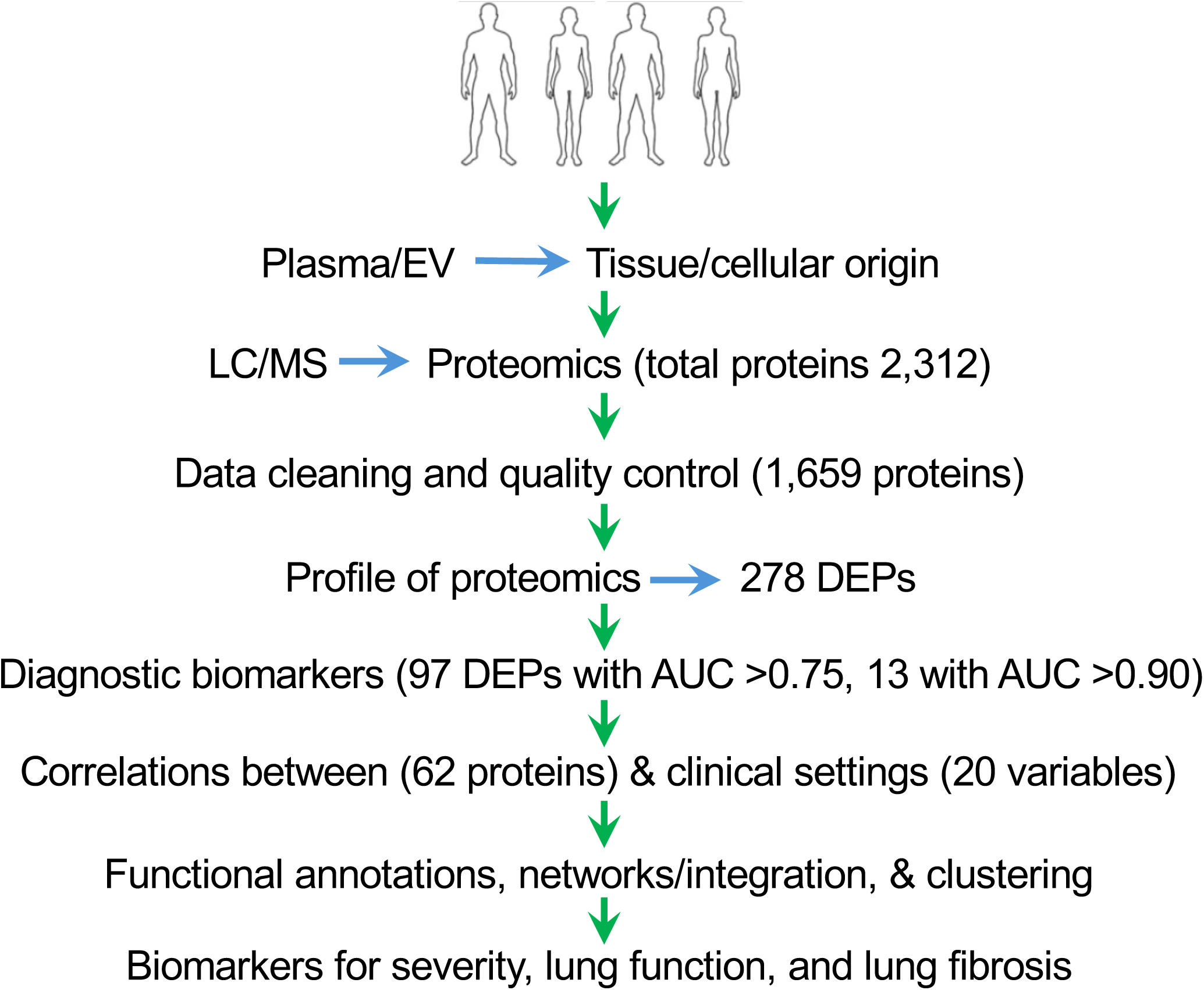
Schematic design of the study. This figure illustrates the various stages of the study, starting from the selection of participants to sample preparation, LC/MS, proteomics data cleaning, profiling, identification of biomarkers, and correlations with clinical settings.

**Table 1.**
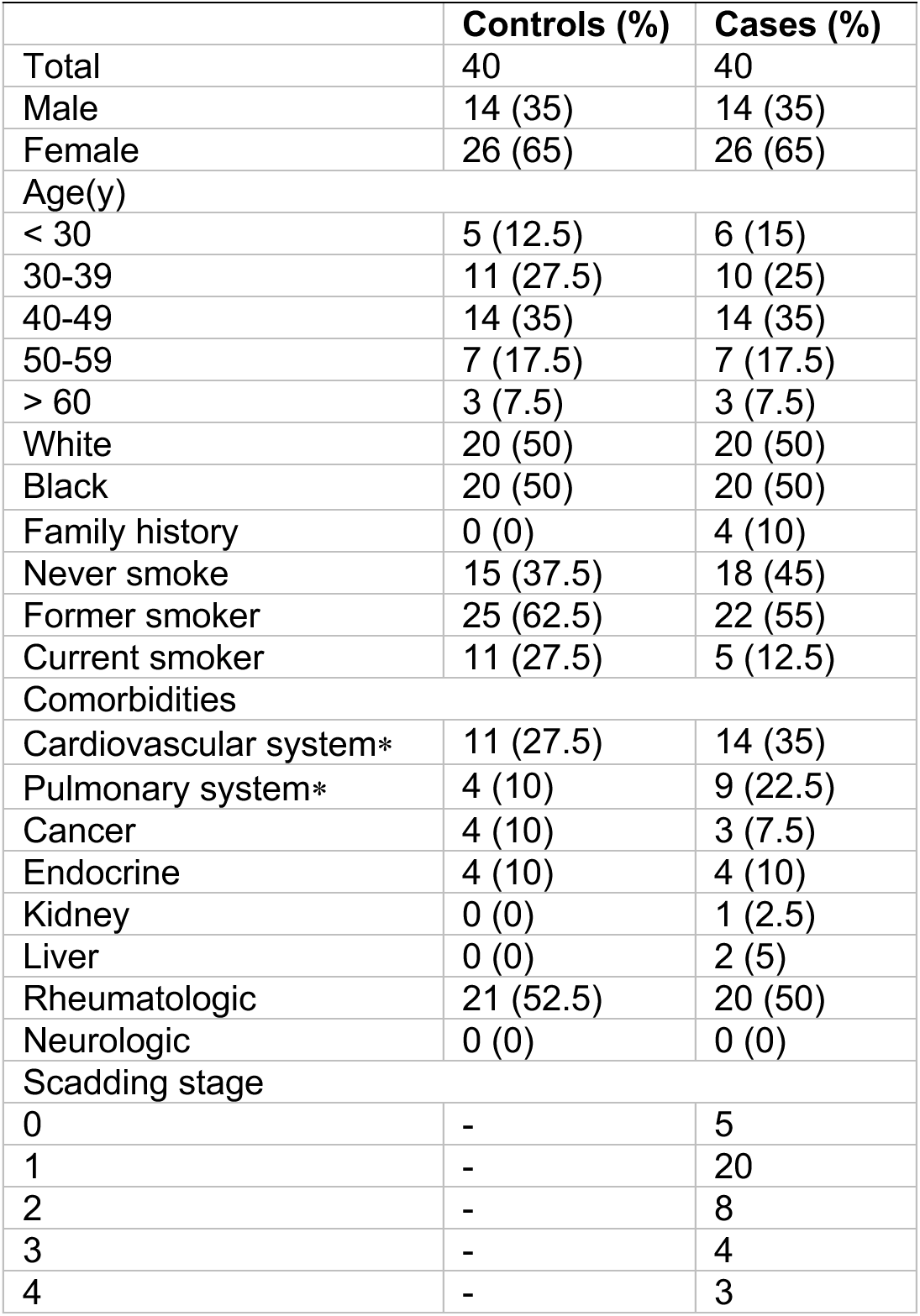
Baseline characteristics of participants.

|  | **Controls (%)** | **Cases (%)** |
| --- | --- | --- |
| Total | 40 | 40 |
| Male | 14 (35) | 14 (35) |
| Female | 26 (65) | 26 (65) |
| Age(y) |  |  |
| < 30 | 5 (12.5) | 6 (15) |
| 30-39 | 11 (27.5) | 10 (25) |
| 40-49 | 14 (35) | 14 (35) |
| 50-59 | 7 (17.5) | 7 (17.5) |
| > 60 | 3 (7.5) | 3 (7.5) |
| White | 20 (50) | 20 (50) |
| Black | 20 (50) | 20 (50) |
| Family history | 0 (0) | 4 (10) |
| Never smoke | 15 (37.5) | 18 (45) |
| Former smoker | 25 (62.5) | 22 (55) |
| Current smoker | 11 (27.5) | 5 (12.5) |
| Comorbidities |  |  |
| Cardiovascular system* | 11 (27.5) | 14 (35) |
| Pulmonary system* | 4 (10) | 9 (22.5) |
| Cancer | 4 (10) | 3 (7.5) |
| Endocrine | 4 (10) | 4 (10) |
| Kidney | 0 (0) | 1 (2.5) |
| Liver | 0 (0) | 2 (5) |
| Rheumatologic | 21 (52.5) | 20 (50) |
| Neurologic | 0 (0) | 0 (0) |
| Scadding stage |  |  |
| 0 | - | 5 |
| 1 | - | 20 |
| 2 | - | 8 |
| 3 | - | 4 |
| 4 | - | 3 |

### 2. Proteomic profile of the discovery cohort

The high-throughput LC-MS pipeline successfully identified 2,312 proteins in the plasma samples. Following the removal of proteins with low abundance, along with data normalization and imputation, a total of 1,659 high-quality proteins remained. Among them, 278 proteins exhibited notable differences in abundance (adjusted p-value < 0.05, |FC| > 1.2), named as differentially expressed proteins (DEPs) (**Figure 2A**). Of these, 118 were upregulated, and 160 were downregulated compared to the controls. The upregulated DEPs were hierarchically clustered into 4 subsets on the heatmap, while the downregulated DEPs were clustered into 6 subgroups. As depicted on the volcano plot (**Figure 2B**), the most downregulated protein in sarcoidosis was calpain-1 catalytic subunit (P07384, log_2_FC, -4.9 and log_10_P, 14.9), followed by acyl-CoA-binding domain-containing protein 6 (Q9BR61, log_2_FC, -3.6 and log_10_P, 6.5). The most increased proteins in sarcoidosis were uncharacterized protein C6orf132 (Q5T0Z8, log_2_FC, 1.7 and -log_10_P, 11) and zinc finger protein 607 (Q96SK3, log_2_FC, 2.5 and -log_10_P, 7.0). The first and second principal components explained approximately 22.3% and 10.4% of the total variance, respectively (**Figure 2C**). The lack of overlap between the two groups indicates that these proteins hold great promise as diagnostic biomarkers to differentiate sarcoidosis patients from healthy controls.

**Figure 2.**
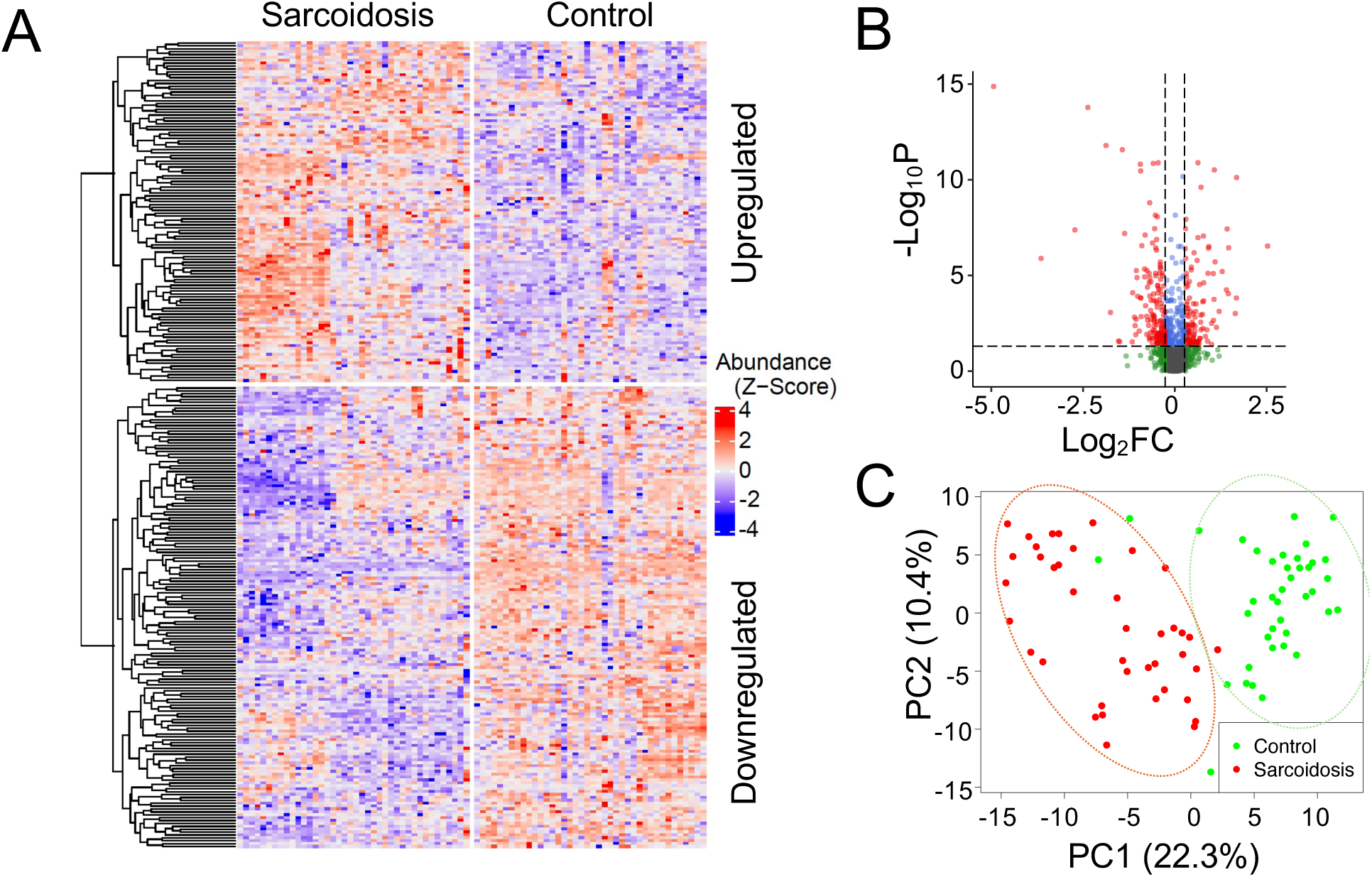
Profile of plasma proteomics in sarcoidosis. **A.** Heatmap. Hierarchical clustering was performed on 278 differentially expressed proteins (DEPs) between controls and patients and visualized in a heatmap. Each row corresponds to one protein. The protein abundance is first preprocessed (see Method) and then standardized to z-scores with mean zero and standard deviation one. Upregulate (top) and downregulated (bottom) proteins were separated. Each column represents one individual in the cohort. **B.** Volcano plot. The proteomics dataset was log-transformed for fold change (FC) on the x-axis (log_2_FC) and for the adjusted p-value on the y-axis (-log_10_P). Horizontal and vertical dashed lines indicate that adjusted p < 0.05 and FC ≥ 1.2 thresholds, respectively. **C.** Principal component analysis (PCA). PCA dimension reduction was performed using the DEPs. The DEPs differentiate controls (green dots) from patients (red dots), as demonstrated within the green and orange ovals, respectively. The first two principal components were displayed on each axis of the plot.

### 3. Tracking tissue and cellular origin

EVs can be released by various tissues and cells into the bloodstream. To track the tissue origin of DEPs in EVs, we searched the TopGene database (**Figure 3**). The top-10 contributing tissues to the identified DEPs were peripheral blood mononuclear cells (PBMC), bronchoalveolar lavage fluid (BALF), epithelium, spleen, lymph node, endothelium, small intestine, airway, lung, and blood. Similarly, the top-ranked cells included platelets, megakaryocytes, dendritic cells, neutrophils, myeloids, club cells, monocytes, CD14^+^ cells, macrophages, and aerocytes. Taken together, the majority of EVs in the plasma were released from inflammatory cells and alveolar epithelial cells, indicating the presence of pulmonary inflammation.

**Figure 3.**
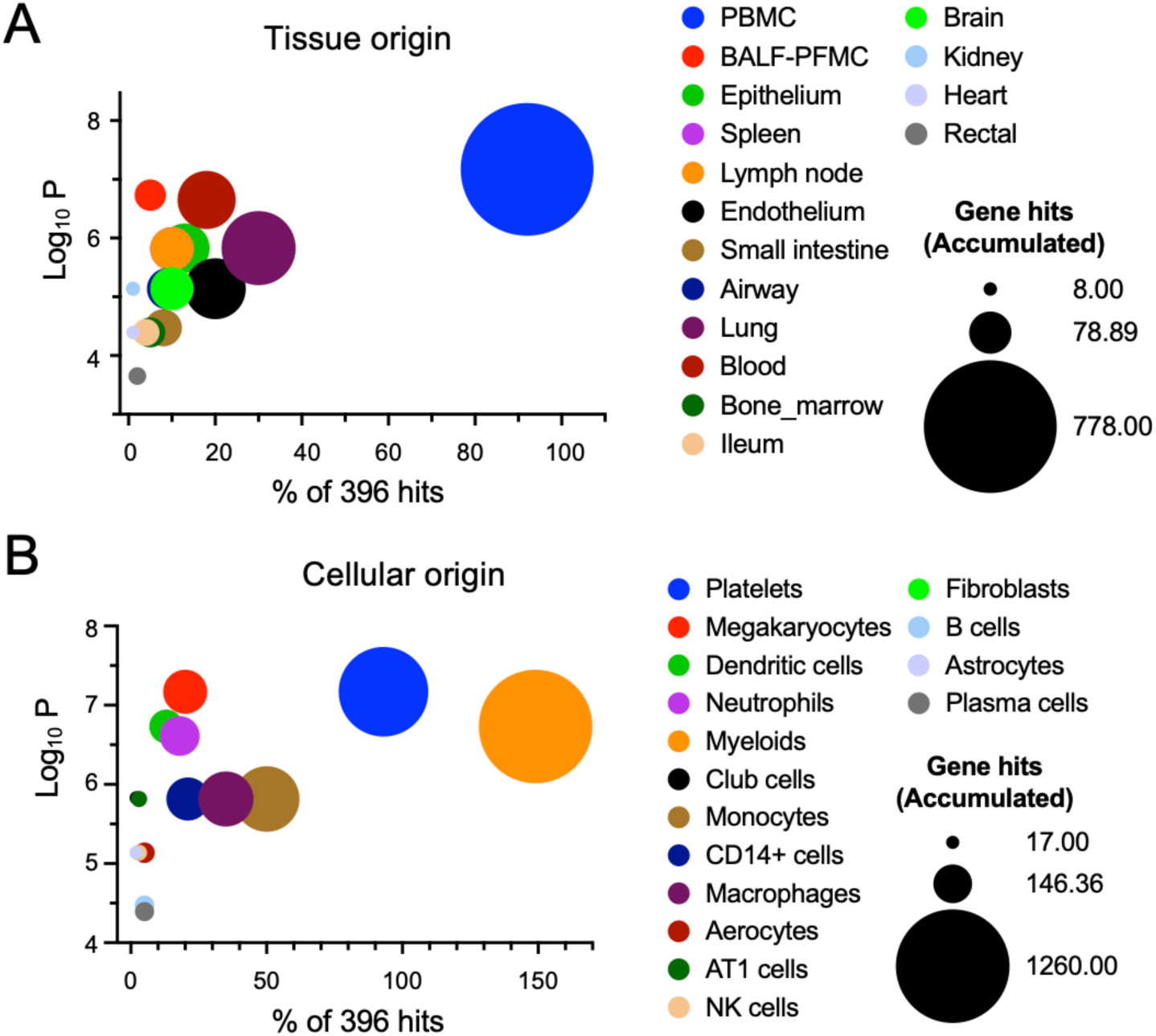
Tissue and cellular origin of exosomal DEPs. The tissue and cellular origin were determined by measuring the frequency of cells and tissues associated with the DEPs using the TopGene databases. **A.** Tissue origin of exosomal proteins. **B.** Cellular origin of exosomal proteins.

### 4. Identification of diagnostic biomarkers for sarcoidosis

To identify EV biomarkers that differentiate sarcoidosis patients from healthy controls, we calculated the area under the curve (AUC) of receiver operating characteristics (ROC) curve for the 278 DEPs individually. The AUC values ranged from 0.583 to 0.971, with 97 proteins having an AUC value above 0.750. Of note, the AUC exceeded 0.900 for the top-13 ranked biomarker candidates (**Table 2**). These 13 biomarkers exhibited high predictive performance for diagnosis, as reflected by their corresponding sensitivity, specificity, accuracy, positive predictive values (PPV), and negative predictive values (NPV). Their functions include anti-infection, host inflammatory responses, cell proliferation, and glucose metabolism. **Figure 4** illustrates the ROC curves of the top-10 candidate biomarkers, including calpain-1 catalytic subunit (P07384, AUC 0.971, sensitivity 100%, specificity 92.5%), probable maltase-glucoamylase 2 (Q2M2H8, 0.962, 95.0%, 92.5%), and synaptotagmin-like protein 4 (Q96C24, 0.947, 92.5%, 95.0%). The protein abundance of these biomarker candidates exhibited a significant difference between patients and controls. Except for toll-like receptor 8 (Q9NR97) and dynein axonemal heavy chain 8 (Q96JB1), the other 8 biomarker proteins, including Calpain-1 catalytic subunit (P07384), Probable maltase-glucoamylase 2 (Q2M2H8), Synaptotagmin-like protein 4 (Q96C24), Protein argonaute-3 (Q9H9G7), Metal-response element-binding transcription factor 2 (Q9Y483), Protein WWC3 (Q9ULE0), Selenoprotein P (P49908), Fukutin related protein (M0QYV8), Uncharacterized protein C6orf132 (Q5T0Z8), Lysozyme C (P61626), and Testin (Q9UGI8), showed a marked reduction (p < 0.0001) in the patient group. Our study suggests that both upregulated and downregulated EV proteins in the plasma could serve as diagnostic biomarkers.

**Figure 4.**
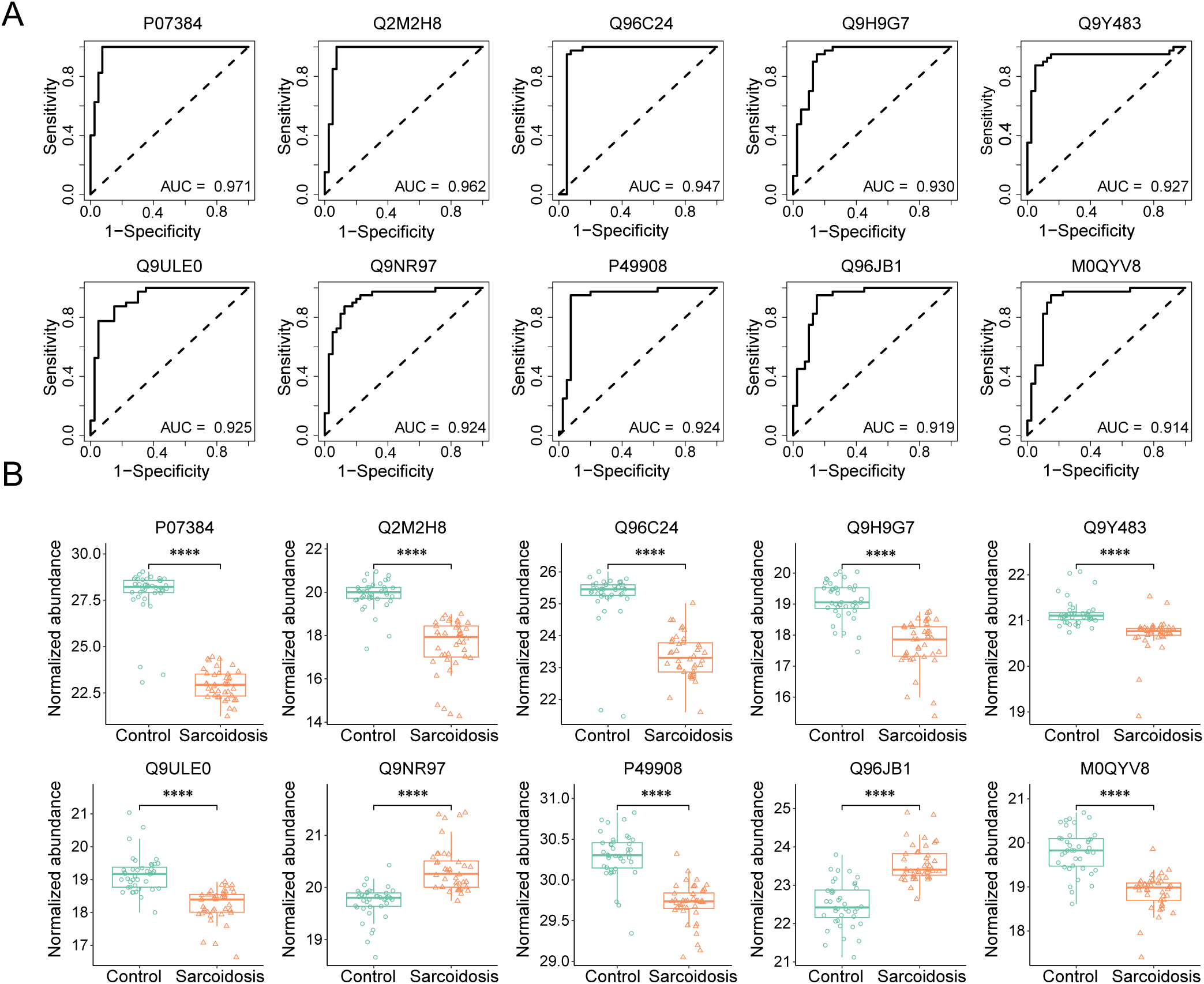
Identification of diagnostic biomarkers and comparison of their expression levels. **A.** Top 10 biomarker candidates with an AUC value >0.900. Performance metrics were averaged over 100 repetitions of the five-fold cross-validation (CV) using univariable logistic regression. The protein biomarkers were graphed in descending order of AUC value from left to right. PPV, positive predictive value. NPV, negative predictive value. **B.** Expression levels of the top-10 identified biomarkers. The box and whisker plots showed the differences in protein abundance between controls and patients. The first quartile, median, third quartile, range, and normalized protein abundance **** adjusted p < 0.0001, Wilcoxon rank-sum test.

**Table 2.** Prediction performance of top 13 potential biomarkers. Area under the curve (AUC), sensitivity (SNS), specificity (SPC), accuracy, PPV (positive predictive value), and NPV (negative predictive value). The information on these proteins, including ID, description, and function, is retrieved from the UniProt database.

| Biomarker | Description | AUC | SNS (%) | SPC (%) | Accuracy | PPV | NPV | Function |
| --- | --- | --- | --- | --- | --- | --- | --- | --- |
| P07384<br>CAPN1 | Calpain-1 catalytic subunit | 0.971 | 100.0 | 92.5 | 0.963 | 0.930 | 1.000 | Ca-regulated non-lysosomal thiol-protease, which catalyzes substrates involved in cytoskeletal remodeling and signal transduction |
| Q2M2H8<br>MGAM2 | Probable maltase-glucoamylase 2 | 0.962 | 95.0 | 92.5 | 0.938 | 0.927 | 0.949 | D-glucose metabolism |
| Q96C24<br>SYTL4 | Synaptotagmin-like protein 4 | 0.947 | 92.5 | 95.0 | 0.938 | 0.949 | 0.927 | Modulates exocytosis of dense-core granules and secretion of hormones in the pancreas and the pituitary |
| Q9H9G7<br>AGO3 | Protein argonaute-3 | 0.930 | 82.5 | 87.5 | 0.850 | 0.868 | 0.833 | Required for RNA-mediated gene silencing (RNAi) |
| Q9Y483<br>MTF2 | Metal-response element-binding transcription | 0.928 | 92.5 | 85.0 | 0.888 | 0.860 | 0.919 | transcriptional networks, methylation activity |
| Q9ULE0<br>WWC3 | Protein WWC3 | 0.925 | 87.5 | 82.5 | 0.850 | 0.833 | 0.868 | Hippo signaling, cell fate |
| Q9NR97<br>TLR8 | Toll-like receptor 8 | 0.924 | 80.0 | 90.0 | 0.850 | 0.889 | 0.818 | innate and adaptive immunity |
| P49908<br>SELENOP | Selenoprotein P | 0.924 | 90.0 | 92.5 | 0.913 | 0.923 | 0.902 | responsible for extracellular antioxidant defense of selenium |
| Q96JB1<br>DNAH8 | Dynein axonemal heavy chain 8 | 0.919 | 95.0 | 82.5 | 0.888 | 0.844 | 0.943 | produces force towards the minus ends of microtubules, sperm motility, ATPase activity |
| M0QYV8<br>RKRP | Fukutin related protein | 0.914 | 92.5 | 85.0 | 0.888 | 0.860 | 0.919 | n/a |
| Q5T0Z8<br>C6orf132 | Uncharacterized protein C6orf132 | 0.904 | 70.0 | 90.0 | 0.900 | 0.882 | 0.783 | n/a |
| P61626<br>LYZ | Lysozyme C | 0.903 | 82.5 | 82.5 | 0.825 | 0.825 | 0.825 | bacteriolytic function of monocytes and macrophages |
| Q9UGI8<br>TES | Testin | 0.900 | 82.5 | 87.5 | 0.875 | 0.868 | 0.833 | cell adhesion, cell spreading, cell proliferation, actin cytoskeleton reorganization, tumor suppression |

### 5. Correlations between biomarker candidates and clinical variables

The clinical assessment of sarcoidosis relies symptoms, chest imaging, lung functional tests, electrophysiological tests, and blood tests. We hypothesize that the identified EV biomarker candidates correlate with these clinical tests. Pearson correlation coefficient was calculated between the protein biomarkers and the following clinical variables: blood tests, chest X-rays, spirometry, radiographic Scadding stage, and dyspnea scale (**Figure 5**). Three proteins, namely Pulmonary surfactant-associated protein B (P07988), DnaJ homolog subfamily A member 2 (O60884), and Immunoglobulin heavy variable 6-1 (A0A0B4J1U7), were found to correlate with the Scadding stage (p < 0.05, |R| > 0.4). Serine/threonine-protein kinase PLK2 (Q9NYY3) correlated with the dyspnea scale. Regarding the lung functional tests, 16 proteins were correlated with 7 clinical variables, including FEV1 (correlated with 3 proteins), FEVPRD (4 proteins), FVC (3 proteins), FVCPRD (8 proteins), PERFEV (1 protein), PERFVC (1 protein), and FEV/FVC ratio (3 proteins). Then, we validated these correlations by analyzing the links between chest X-rays and biomarker candidates. In total, there were 39 proteins correlated with 6 X-ray readouts (p < 0.05, |FC| > 1.4). These readouts included lymphadenopathy (corrected with 7 proteins), alveolar infiltrates (9 proteins), parenchymal bulla or blebs (4 proteins), hilar retraction (17 proteins), interstitial infiltrate (1 protein), and importantly, pulmonary fibrosis (3 proteins). Third, blood tests were correlated with 8 proteins (p < 0.05, |R| > 0.5. These blood tests were creatinine level (4 proteins), WBC (1 protein), eosinophils (1 protein), potassium content (1 protein), and blood urea nitrogen (BUN, 1 protein). In total, 54 EV proteins were identified that correlated with the analyzed clinical variables (**Figure 5D**). Among them, 8 proteins were shared by the three subgroups of clinical variables. More than half of the correlated proteins (29 out of 54) were identified as potential biomarkers (AUC > 0.750 as a broadly used cutoff criterion), with 2 of them, namely, protein WWC3 (Q9ULE0) and uncharacterized protein C6orf132 (Q5T0Z8) being top-ranked candidates (AUC > 0.90). These two proteins were correlated with alveolar infiltrates and hilar retraction, respectively. Furthermore, the correlations between the selected 10 biomarkers (AUC > 0.750) and continuous clinical variables were visualized using linear regression (**Figure 6A**). These 9 proteins were serine/threonine-protein kinase PLK2 (Q9NYY3), small ribosomal subunit protein eS8 (P62241), kinesin-1 heavy chain (P33176), chitinase-3-like protein 1 (P36222), peptidyl-prolyl cis-trans isomerase D (Q08752), DNAJ homolog subfamily A member 2 (O60884), cytoplasmic tryptophan--tRNA ligase (P23381), thrombospondin-4 (P35443), and uncharacterized protein C16orf46 (Q6P387). Two-thirds of the correlations were negative, while one-third were positive. Among the correlations, the slope of 6 plots was above 1.0 and up to 7.0. These results suggest a strong link between differential proteins and clinical variables, suggesting these EV proteins could function as clinical biomarkers.

**Figure 5.**
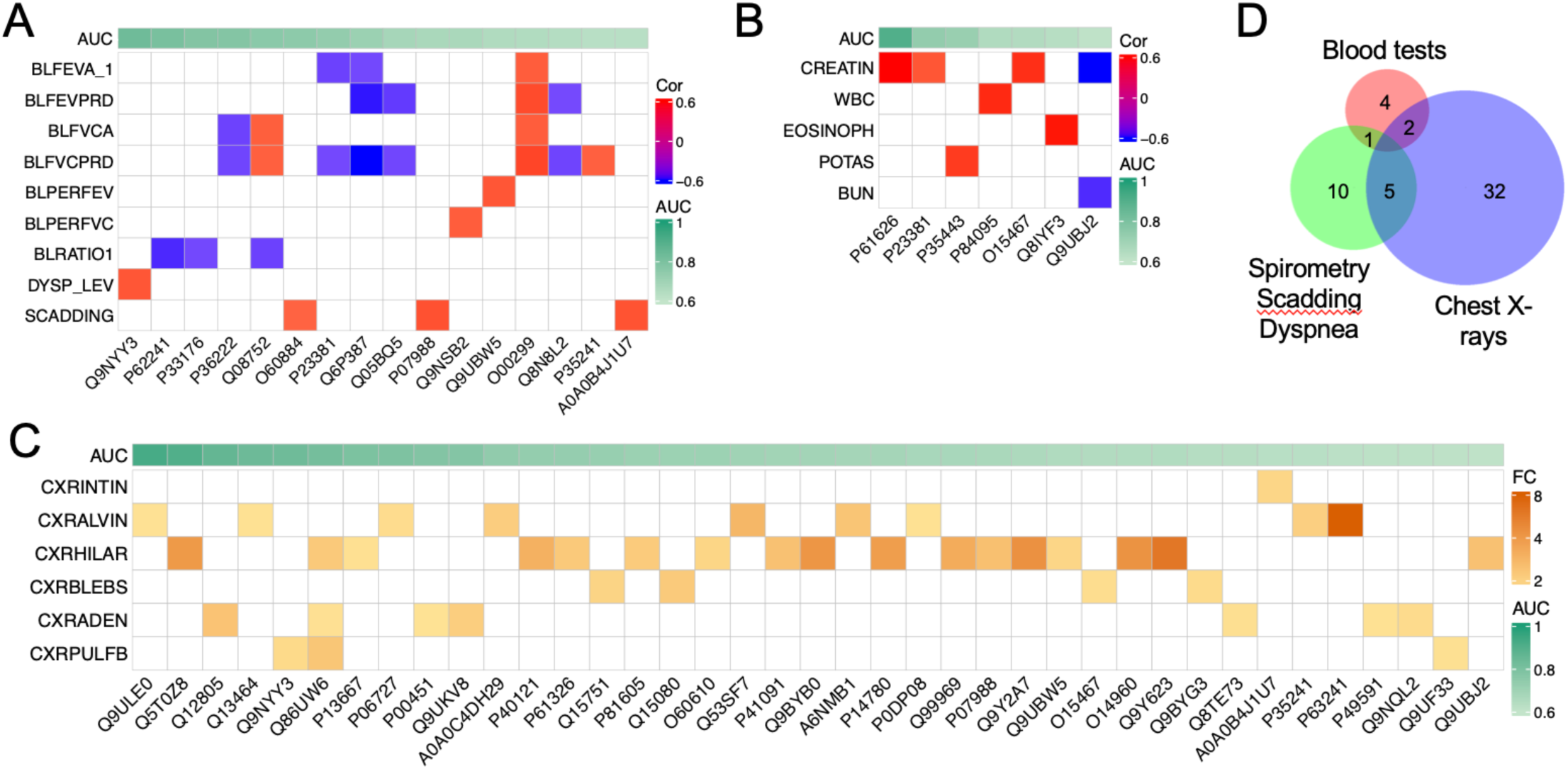
Correlations between identified biomarkers and three subgroups of clinical variables. Correlations between biomarker candidates and clinical variables were sorted by their AUC values. **A.** Correlation matrix between spirometry and biomarker candidates. Sixteen proteins correlated with 7 spirometry variables, Scadding stage (SCADDING), and dyspnea scale (DYSP_LEV). Seven spirometry variables were pre-bronchodilators FEV-1 (BLFEVA_1), FEV-1 predicted (BLFEVPRD), pre-bronchodilators FVC (BLFVCA), FVC predicted (BLFVCPRD), FEV-1 % predicted (BLPERFEV), FVC-1 % predicted (BLPERFVC), FEV-1/FVC (BLRATIO1). p-value < 0.05, Pearson correlation coefficient > 0.4 (shown by colors). **B.** Correlations between blood tests and biomarker candidates. Seven proteins correlated with 5 variables of blood tests. The correlated blood tests included creatinine level (CREATIN, mg/dL), white blood cells (WBC, x10^3^/mm^3^), eosinophils (EOSINOPH, %), plasma potassium (POTAS, mEq/L), and blood urea nitrogen level (BUN, mg/dL). P-value < 0.005, Pearson correlation coefficient > 0.5. **C.** Correlation matrix between biomarker candidates and chest X-ray readouts. Six image readouts and 39 biomarker candidates were correlated. The six x-ray readouts were interstitial infiltrates (CXRINTIN), alveolar infiltrates (CXRALVIN), Hilar retraction (CXRHILAR), Bullae or blebs (CXRBLEBS), adenopathy (CXRADEN), and pulmonary fibrosis (CXRPLUFB). P-values < 0.05, FC > 2.0 for categorical variables. **D.** Venn plot. Unique and shared biomarker proteins correlated to the three subgroups of clinical variables are illustrated.

**Figure 6.**
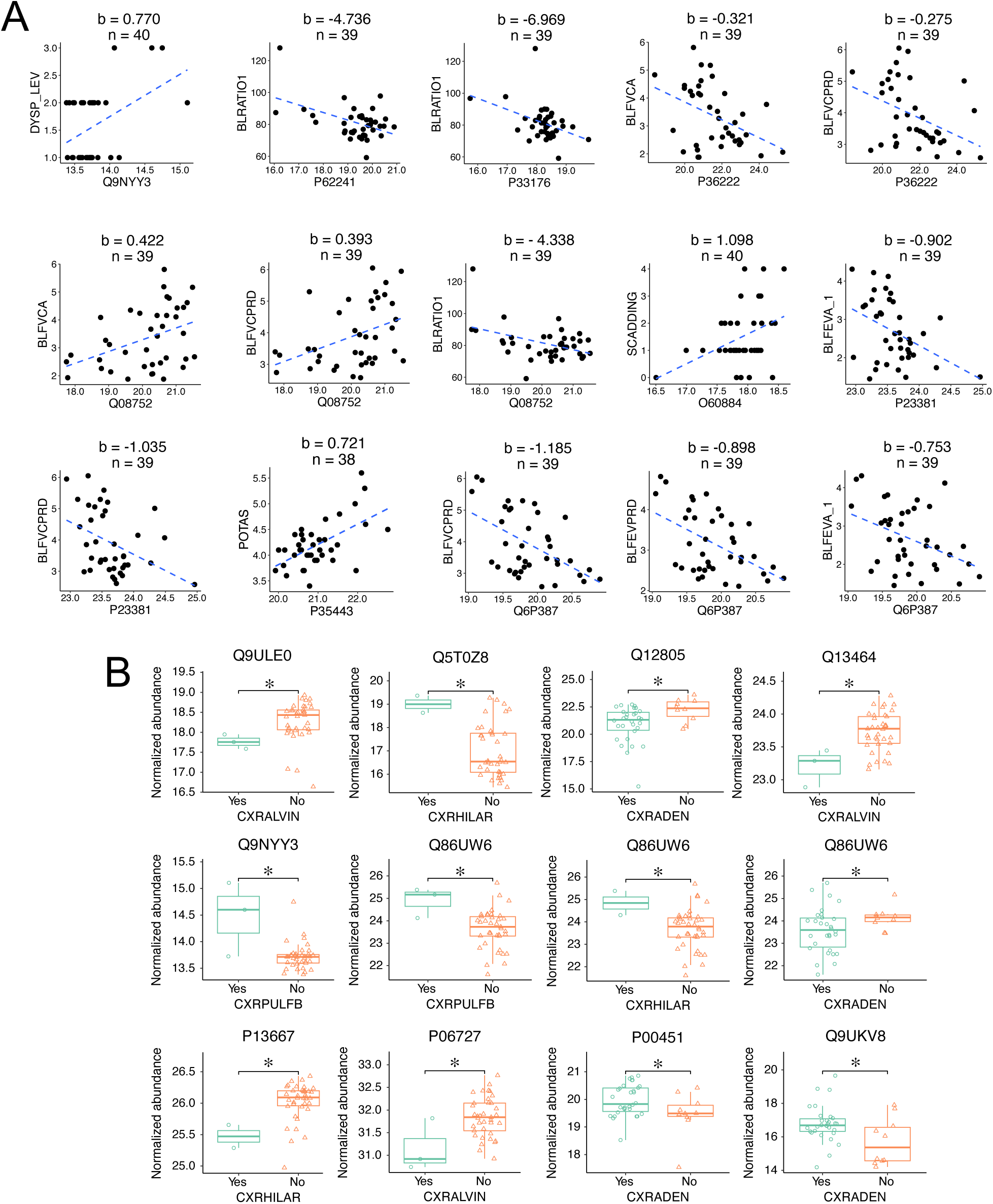
Linear and binary correlations between biomarker candidates and the top-correlated clinical variables. **A.** Linear regression of the top-ranked continuous variables. The plots were sorted by their AUC values descendingly. Clinical variables were plotted as a function of normalized protein abundance. The blue dashed lines were generated by regressing the clinical variables with the corresponding normalized protein abundance. The slope (b) and sample size (n) were on the top of the plots. The correlated clinical variables included dyspnea scale (DYSP_LEV), FEV-1/FVC (BLRATIO1), pre-bronchodilators FVC (BLFVCA), FVC predicted (BLFVCPRD), Scadding stage (SCADDING), pre-bronchodilators FEV-1 (BLFEVA_1), and potassium content (POTAS, mEq/L). **B.** Boxplots of protein abundance between the binary outcomes of the top-ranked chest X-ray variables. The plots were sorted by their AUC values descendingly. The normalized protein abundance was compared between negative (No, orange triangle) and positive patients (Yes, aqua circle) for the correlated variables. The boxplot shows the first quartile, median, third quartile, range, and normalized protein abundance. The X-ray readouts were alveolar infiltrates (CXRALVIN), Hilar retraction (CXRHILAR), adenopathy (CXRADEN), and pulmonary fibrosis (CXRPLUFB). * p < 0.05, Wilcoxon rank-sum test.

### 6. Different expression levels of biomarkers correlated with X-ray readouts

Abnormal X-ray readouts were correlated with specific biomarkers. Notably, a significant difference existed in the expression levels of correlated protein biomarkers between patients with and without abnormal X-ray findings (**Figure 6B**). These 10 biomarker proteins exhibited an AUC value > 0.774, namely, protein WWC3 (Q9ULE0), uncharacterized protein C6orf132 (Q5T0Z8), EGF-containing fibulin-like extracellular matrix protein 1 (Q12805), Rho-associated protein kinase 1 (Q13464), serine/threonine-protein kinase PLK2 (Q9NYY3), NEDD4-binding protein 2 (Q86UW6), protein disulfide-isomerase A4 (P13667), apolipoprotein A-IV (P06727), coagulation factor VIII (P00451), and protein argonaute-2 (Q9UKV8). Half of these biomarker proteins were upregulated in patients with abnormal chest X-ray reports: 2 for pulmonary fibrosis, 2 for hilar retraction, and 1 for lymphadenopathy. In comparison, the expression of 6 biomarker proteins was downregulated in patients with alveolar infiltrates ((protein WWC3 (Q9ULE0), Rho-associated protein kinase 1 (Q13464), and apolipoprotein A-IV (P06727)), lymphadenopathy ((EGF-containing fibulin-like extracellular matrix protein 1 (Q12805) and NEDD4-binding protein 2 (Q86UW6)), and hilar retraction (protein disulfide-isomerase A4, P13667). Protein Q86UW6 was correlated with lung fibrosis, hilar retraction, and lymphadenopathy. Pulmonary fibrosis showed increased expression of serine/threonine-protein kinase PLK2 (Q9NYY3) and NEDD4-binding protein 2 (Q86UW6), while alveolar infiltrates exhibited reduced expression of protein WWC3 (Q9ULE0), Rho-associated protein kinase 1 (Q13464), and apolipoprotein A-IV (P06727). These biomarkers hold the potential as predictors of the clinical course of pulmonary sarcoidosis.

### 7. Functions and pathway enrichment networks of biomarker candidates

To enrich the functions and network of the biomarker candidates with an AUC > 0.750, hierarchical clustering of the functional pathways, visualization of top-ranked biomarker candidates, and network analysis were performed (**Figure 7**. The top three pathways were clathrin-mediated endocytosis, Hsp90 chaperone cycle for steroid hormone receptors in the presence of ligand, and spliceosome (**Figure 7A**). The top 30 pathways could be grouped into three subgroups: endocytosis, responses to external stimuli, and transcription, which were enriched by 9, 13, and 5 biomarker proteins, respectively (**Figure 7B**). Moreover, both upregulated and downregulated biomarker proteins were analyzed for their network interactions (**Figure 7C**). There were 2 pathways in the group for endocytosis: clathrin-mediated endocytosis and endocytosis, 6 pathways in the group of response to external stimuli, and 2 in the group for transcription. Interactions of the “endocytosis” and “response to external stimuli” were mediated by 4 biomarker proteins, namely, F-actin-capping protein subunit alpha-2 (P47755), HLA class I histocompatibility antigen B alpha chain (D9J307), HLA class I antigen (Q53Z42), and E3 ubiquitin-protein ligase CBL (A0A0U1RQX8). These proteins were shared by 2 endocytosis pathways and 3 pathways belonging to the group of “response to external stimuli": Hsp90 chaperone cycle, antigen processing and presentation, and the VEGFR1 pathway. In contrast, 2 pathways in the “transcription” group did not interact with each other or with the pathways from the other two groups. The functional enrichment and network analysis suggest that clathrin-mediated endocytosis, Hsp90, and spliceosome could play a major role in the development of sarcoidosis.

**Figure 7.**
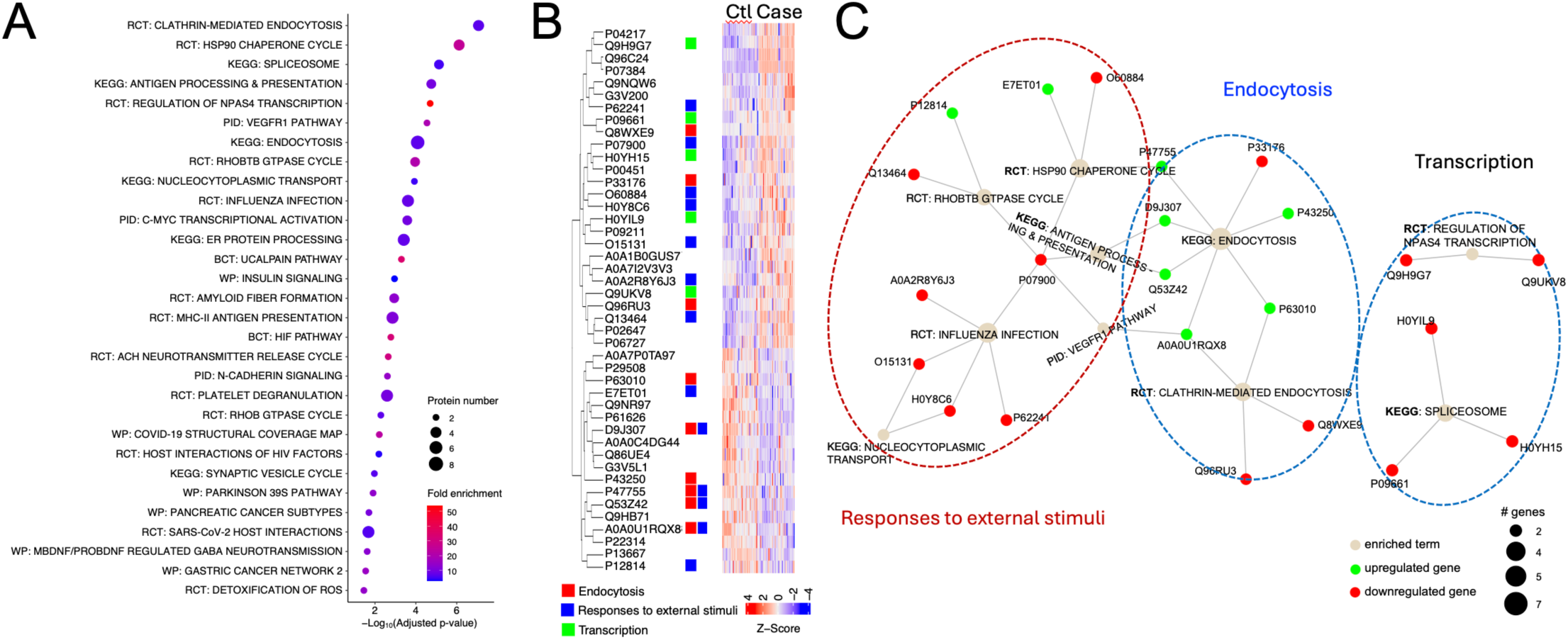
Functional annotations and network analysis. **A**. Dotplot of enriched pathways of biomarkers with an AUC value > 0.750. The top 30 clustered signaling pathways were plotted. Dot size indicates the number of proteins and colors represent fold enrichment. Pathways were ordered by the significance level of the representative pathway terms. RCT, Reactome; KEGG, Kyoto Encyclopedia of Genes and Genomes, PID, Pathway Interaction Database; BCT, Biocarta; WP, Wikipathways. **B.** Integrated functions of biomarker candidates in the top 30 pathways shown in panel **A**. The Z-score of the heatmap showed the relative expression levels of the proteins that were top-ranked based on their AUC values for both controls (Ctl) and sarcoidosis (Case). Three groups of pathways and corresponding biomarkers were marked with red, blue, and green color for endocytosis, responses to external stimuli, and transcription, respectively. **C**. Integrated networks of the top 10 enriched pathways shown in panel **B**. Proteins were connected with their respective pathways.

## Discussion

The utilization of extracellular vesicles (EVs) in conjunction with proteomics facilitates the identification of potential diagnostic plasma biomarkers for sarcoidosis. Our application of EV proteomics represents an innovative method for the discovery of novel biomarkers. In addition, these biomarkers validated previously identified biomarkers of disease severity in sarcoidosis. The tissue and cellular origins of plasma EV proteins are systemic. The identified EV biomarkers exhibit a strong correlation with essential clinical variables related to disease severity, laboratory findings, pulmonary function test findings, and the severity of dyspnea. The signaling pathways and networks associated with the identified EV biomarkers provide insights into the pathogenesis of sarcoidosis. This study represents the initial demonstration of the efficacy of EV omics as a high-throughput approach for biomarker discovery in sarcoidosis.

Seven of the 278 biomarkers identified in this study have been previously reported. Among the top 13 biomarkers with an AUC > 0.90, plasma lysozyme (P61626, encoded by the LYZ gene) and toll-like receptor 8 (Q9NR97, encoded by the TLR8 gene) have been identified as diagnostic biomarkers specifically for pulmonary sarcoidosis ^55^ and cardiac sarcoidosis ^56^, respectively. In addition to the aforementioned top 13 biomarkers, a previous study identified AP2B1 as a potential biomarker for sarcoidosis through proteomic analysis of alveolar macrophages ^57^. In addition, membrane-associated HGFA, ILF3, RAB7A, and RPL18 in alveolar macrophages were proposed as diagnostic biomarker candidates for sarcoidosis ^58,59^. Numerous proteins that we identified have been previously associated with sarcoidosis. Apolipoprotein A1 (P02647) could be a biomarker that has been associated with chronic obstructive pulmonary disease (COPD), tuberculosis (TB), and sarcoidosis^60^. Complement C5 (P01031) and C8a (P07357) are actively synthesized in alveolar macrophages in sarcoidosis ^61^. Whole exome sequencing identified that the protein DNAH11 may contribute to the formation of the characteristic lesion through regulating G-proteins in pediatric sarcoidosis ^62^. Moreover, GSTP1 has been recognized as a central node in the functional enrichment analysis of proteomics in pulmonary sarcoidosis ^60^. The serum concentration of C-C motif chemokine ligand 18 (CCL18) was observed to be markedly increased in individuals diagnosed with intrathoracic sarcoidosis. In contrast, the detection of increased levels of CCL16 (O1546) in the bronchoalveolar lavage (BAL) of individuals diagnosed with sarcoidosis has shown inconsistent results ^63,64^.

We found evidence that some of our EV biomarkers correlate with lung fibrosis. The identified EV biomarkers CCL18 could be implicated in the prediction of sarcoidosis-associated fibrosis. CCL18 has been suggested as a marker for identifying patients at a higher risk of developing pulmonary fibrosis or progressive disease ^65^.

However, CCL18 also serves as a biomarker for interstitial lung disease (ILD) due to its elevated levels in serum, bronchoalveolar lavage (BAL), and alveolar macrophages ^66^. Of note, significant myofibrosis in neuromuscular sarcoidosis is correlated with increased expression of CCL18 in M2 macrophage phenotype ^67^. These studies suggest that some of the EV biomarkers could be shared with other fibrotic diseases. We found that serine/threonine-protein kinase PLK2 (Q9NYY3) protein level in plasma EVs was highly correlated with the occurrence of lung fibrosis in sarcoidosis. Genetic knockout of the corresponding gene PLK2 leads to the manifestation of a lung fibrosis phenotype ^68^. The second lung fibrosis-correlated EV protein, Q9UF33 (encoded by the EPHA6 gene), is implicated in the severity of bleomycin-induced pulmonary fibrosis in mice ^69^. Moreover, EPHA3^+^ lung cells from IPF patients induce lung fibrosis in mice, and antibody-mediated depletion of these cells ameliorates fibrosis ^70^.

Several of the top thirty-ranked signaling pathways enriched by the identified EV biomarkers have been reported previously. The upregulation of two phagocytotic pathways, namely Fcγ receptor-mediated phagocytosis and clathrin-mediated endocytosis, has been documented in alveolar macrophages in sarcoidosis ^57,59^. These pathways are associated with the functionality of alveolar macrophages. A robust correlation has been reported between HSP-70 and uveitis in patients diagnosed with sarcoidosis ^71^. Furthermore, the expression level of adenosine diphosphate-ribosylation factor GTPase activating protein 1 is notably elevated in sarcoidosis compared to asthma ^72^.

There are some limitations to this study. Although the plasma samples were collected by a multi-centered clinical trial -ACCESS, all these patients were from the United States and may not be representative of world-wide sarcoidosis. An independent validation cohort could be applied to confirm the identified EV biomarkers. This is a single omics study based on the analysis of plasma EV proteomes. Other omics datasets from the plasma and other liquid samples (e.g., urine, BAL), including plasma EV RNA profiling, EV DNA analysis, and EV metabolomics, could improve the accuracy of the biomarkers identified in this study. Long-term storage of the ACCESS samples could be a concern. However, the considerable stability of EV proteins compensates for the alterations associated with long-term storage.

In summary, this study demonstrates that hundreds of EV biomarkers can be identified by combining unbiased high-throughput proteomics and machine-learning algorithms. , Many of the biomarkers that we identified were associated with previously known clinical findings, biomarkers and immunologic pathways that have been thought to be important in sarcoidosis, This suggests that our discovered biomarkers have great clinical relevance.Our results provide novel biomarker candidates for designing prospective clinical trials to identify biomarkers for phenotype-specific, organ-specific, predictive, and treatment responses in sarcoidosis.

## Author contributions

HLJ conceived and designed the study. HLJ, NMX, CM, MAJ, LLK, and RZ contributed to the writing of the manuscript. NMX, LG, EG, CM, MQ, DZ, and HLJ analyzed the data and performed the statistical analysis. LLK, JWZ, and MAJ reviewed the manuscript and provided critical comments.

## Acknowledgments

This work was supported by NHLBI: grant NIH grant HL87017 (to HLJ) and institutional funds (to HLJ).

## Ethics approval and consent to participate

All participants provided written informed consent through the ACCESS clinical trial. This study was approved by the Institutional Review Board (LU#216964) at Loyola University Chicago.

## Competing financial interests

None of the authors of this manuscript had any financial relationship with a commercial company except MAJ who received grants for his institution from Xentria Pharmaceuticlas and aTyr Pharmaceuticals and is a consultant for Merck, Priovant Pharmaceuticals, and Sparrow Pharmaceuticals.

### Abbreviation list

ACE: angiotensin-converting enzyme
ACCESS: A Case Controlled Etiologic Study of Sarcoidosis
AUC: area under the curve
BALF: bronchoalveolar lavage fluid
BUN: blood urea nitrogen
CRP: C-reactive protein
CTSS: cathepsin S
DEP: differentially expressed proteins
EGF: epidermal growth factor
EV: extracellular vesicles
FEV1: forced expiratory volume in 1 second
FEVPRD: FEV1% predicted
FVC: forced vital capacity
FVCPRD: forced vital capacity predicted
LBP: lipopolysaccharide-binding protein
LC-MS: liquid chromatography–mass spectrometry
MMP14: matrix metallopeptidase 14
NPV: negative predictive value
PERFEV: FEV-1 % predicted
PERFVC: FVC % predicted
PLK2: polo-like kinase 2
PPV: positive predictive value
ROC: receiver operating characteristics

